# PXDesign: Fast, Modular, and Accurate De Novo Design of Protein Binders

**DOI:** 10.1101/2025.08.15.670450

**Authors:** Protenix Team, Milong Ren, Jinyuan Sun, Jiaqi Guan, Cong Liu, Chengyue Gong, Yuzhe Wang, Lan Wang, Qixu Cai, Wenzhi Ma, Yuxuan Zhang, Zhenyu Liu, Hanyu Zhang, Xinshi Chen, Wenzhi Xiao

## Abstract

PXDesign achieves nanomolar binder hit rates of 17–82% across six of seven diverse protein targets, surpassing prior methods such as AlphaProteo. This experimental success rate is enabled by advances in both binder generation and filtering. We develop both a diffusion-based generative model (PXDesign-d) and a hallucination-based approach (PXDesign-h), each showing strong *in silico* performance that outperforms existing models. Beyond generation, we systematically analyze confidence-based filtering and ranking strategies from multiple structure predictors, comparing their accuracy, efficiency, and complementarity on datasets spanning *de novo* binders and mutagenesis. Finally, we validate the full design process experimentally, achieving high hit rates and multiple nanomolar binders.

To support future research and broaden community adoption, we release the full PXDesign pipeline (https://github.com/bytedance/PXDesign), provide public access to PXDesign through a dedicated web server (https://protenix-server.com), and make all designed binder sequences available at the project page (https://protenix.github.io/pxdesign).

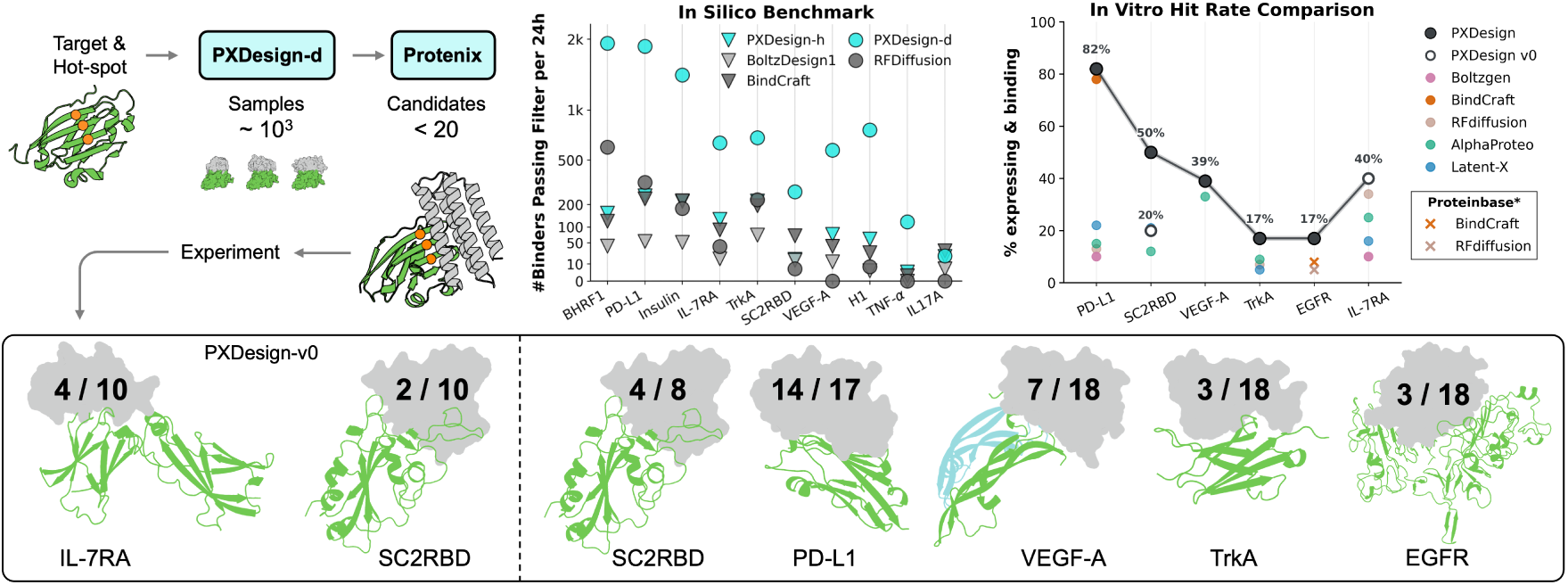

## 1 Introduction

PXDesign is a model suite for de novo protein-binder design that has been validated on seven targets, achieving nanomolar binder hit rates of 17–82% on six of them and outperforming strong baselines such as AlphaProteo [59] by 2–6×. These results reflect a systematic approach targeting two essential aspects of binder design success: proposing structurally complementary candidates, and prioritizing those most likely to bind effectively.

While recent methods have advanced both stages, important gaps remain. Diffusion-based generators [24, 28, 54] and hallucination-based optimization methods [3, 13, 20, 35, 38, 39, 55] have shown promise in proposing viable binder candidates, but direct head-to-head comparisons under consistent evaluation are lacking. Likewise, confidence scores such as pLDDT or interface pTM from the AlphaFold series and related variants [1, 2, 14, 19, 27, 30, 32, 32, 40, 47, 56, 57] are widely used for filtering, but important questions remain: How accurate and generalizable are these scores? Can alternative predictors improve enrichment, diversity, and yield? Addressing these questions is essential to building a robust, high-hit-rate method.

Here, we present PXDesign, a unified framework integrating both binder generation and confidence-based filtering. Our main contributions are:

- **Generation via two strategies**: We develop both a diffusion-based (PXDesign-d) and a hallucination-based (PXDesign-h) generation method, with architectural and algorithmic enhancements for scalability and efficiency. Each achieves state-of-the-art *in silico* performance, and our head-to-head comparison reveals their respective strengths. We also show our diffusion model’s strong capacity for diverse, designable protein generation in unconditional tasks.
- **Filtering strategies**: We construct and evaluate confidence-based filters from Protenix [12] and AlphaFold-based models [1, 27] using datasets including Cao data [11], RFdiffusion [54], EGFR [16] and SKEMPI [25, 34, 36]. Our results highlight the value of alternative predictors and show how filtering strategies influence diversity and yield in complementary ways.
- **Experimental validation across targets**: We evaluated PXDesign on seven diverse protein targets via *in vitro* expression and binding assays. PXDesign achieves high hit rates across six targets (Table 1). Notably, a substantial improvement over the initial version (PXDesign v0) is observed on SC2RBD, where the hit rate increased from 2 out of 10 to 4 out of 8. PXDesign outperformed AlphaProteo [59] across all evaluated targets, achieving 2- to 6-fold higher hit rates on IL-7RA, PD-L1, VEGF-A, SC2RBD, and TrkA.

**Table 1.** Experimental hit rates (%) across different targets and methods. “v0” refers to results from the initial PXDesign version. Results marked with * are sourced from Proteinbase (https://proteinbase.com) and may not fully reflect true performance; all other values are taken from authors’ reports. See Appendix E for details.

| | IL-7RA | PD-L1 | VEGF-A | SC2RBD | TrkA | EGFR | TNF-α |
| --- | --- | --- | --- | --- | --- | --- | --- |
| PXDesign | – | **82** | **39** | **50** | **17** | **17** | 0 |
| PXDesign v0 | **40** | – | – | 20 | – | – | – |
| Boltzgen [45] | 10 | 10 | – | – | – | – | 0 |
| BindCraft [38] | – | 78 | – | – | – | 8* | – |
| RFdiffusion [54] | 34 | 13 | – | – | 7 | 5* | – |
| AlphaProteo [59] | 25 | 15 | 33 | 12 | 9 | – | 0 |
| Chai-1d [48] | 10 | 5 | – | – | – | – | – |
| Chai-2 [48] | 70 | 72 | – | – | – | – | 23 |
| Latent-X ($K_D$ measured) [9] | 16 | 22 | – | 21 | 5 | – | – |
| Latent-X (detectable only) [9] | 26 | 49 | – | 52 | 10 | – | – |

Altogether, PXDesign delivers both high *in vitro* success rates and methodological insights, guiding future applications and enabling reproducible evaluation through open-sourced tools and a public webserver.

## 2 Filtering and Ranking: Accuracy, Efficiency, and Diversity

Filtering and ranking are essential for turning large sets of generated designs into a focused shortlist of promising candidates. In this section, we evaluate **AF2-IG** [54] and **Protenix** variants (full, Mini, and Mini-Templ) [23] across multiple datasets to answer three key questions:

1. **Accuracy**: Which filters enrich true binders most effectively?
2. **Efficiency**: Can we reduce computational cost without sacrificing positive predictive value?
3. **Diversity**: Do different filters capture complementary regions of design space?

### 2.1 Filter Design and Setup

We constructed confidence-based filters using per-sequence and interface-level scores from AF2-IG and Protenix variants. All Protenix models use a 2-step ODE diffusion sampler for efficiency [23]. Thresholds were tuned via grid search on Cao data [11], a retrospective benchmark of experimentally tested designs without AlphaFold-based pre-filtering. The search process is described in Appendix A, and Table 2 summarizes the resulted thresholds.

**Table 2.** Thresholds for combined filters. Each filter represents a model-specific combination of score thresholds used for binary selection. “AF2-IG-easy” reflects thresholds proposed by BindCraft [38]. “AF2-IG” denotes thresholds selected via our own grid search on Cao data; these match the values independently reported in Zambaldi et al. [59]. “AF3” refers to thresholds grid-searched in Zambaldi et al. [59]. “Protenix” denotes a unified threshold set applied to Protenix, Protenix-Mini, and Protenix-Mini-Templ, while “Protenix-basic” represents a relaxed criterion used on challenging targets (e.g., TNF-*α*) in wet-lab experiments to preserve diversity.

| Filter Name | Confidence Thresholds | Structure Thresholds |
| --- | --- | --- |
| AF3 [59] | min ipAE < 1.5, binder pTM > 0.8 | complex RMSD < 2.5 Å |
| AF2-IG [59] | ipAE < 7.0, pLDDT > 0.9 | binder RMSD < 1.5 Å |
| AF2-IG-easy [38] | ipAE < 10.85, ipTM > 0.5, pLDDT > 0.8 | binder bound/unbound RMSD < 3.5 Å |
| Protenix [12] | binder ipTM > 0.85, binder pTM > 0.88 | complex RMSD < 2.5 Å |
| Protenix-basic [12] | binder ipTM > 0.8, binder pTM > 0.8 | complex RMSD < 2.5 Å |

We also tested ranking and enrichment performance on three additional datasets:

- EGFR challenge [15]: expression and binding phenotypes for de novo binders from Adaptyv Bio.
- SKEMPI subset [25]: point mutations with measured ddG, curated to retain high-confidence AF3 structures by Lu et al. [34].
- RFdiffusion wet-lab set [54]: binary binding outcomes for published designs.

### 2.2 Main Findings

**(1) Accuracy — Protenix achieves higher binder enrichment.** On Cao data, Protenix-derived confidence metrics outperform AF2-IG across most targets when used individually (Figure 1a) or in combination (Figure 1b). AF2-IG performs better on a few specific targets, but overall Protenix’s precision and AUC are higher (see Appendix A). On the EGFR challenge, Protenix scores correlate more strongly with experimental expression and binding affinity than AF2 or ESM [30, 44] (Figure 1e). On the SKEMPI subset, Protenix matches or exceeds AF3’s performance, especially in the subset of positive binding ddG mutations (Figure 1f). Importantly, Protenix-based filtering can also improve hit rates on previously published designs. Similarly, in re-ranking the 95 RFdiffusion designs [54], replacing AF2-IG with Protenix ipTM for top 10 or 15 selection substantially increases observed success rates (Figure 1d).
**(2) Efficiency — strong performance at lower cost.** Protenix-Mini and Protenix-Mini-Templ achieve significant runtime reductions compared to the Protenix full model while maintaining comparable enrichment quality, enabling practical large-scale screening and rapid triaging of candidate pools.
**(3) Diversity — complementary coverage of design space.** On Cao data, the overlap between sequences retained by Protenix and AF2-IG is surprisingly small (Figure 1c). Each model captures different subsets of true positives, suggesting that their inductive biases are complementary. This motivates the use of multiple predictors to improve coverage and robustness in real-world applications.

## 3 Generators and In silico Benchmarking

Among various generative strategies explored for protein binder design, two have emerged as the most widely adopted and experimentally validated:

- **Diffusion (training-based):** Generative models, typically denoising diffusion models, trained to sample binder structures conditioned on the target. These approaches require either fine-tuning or training from scratch. Examples include RFdiffusion [28, 54], GeoFlow-v2 [46], Chai-2 [48], Latent-X [9], etc.
- **Hallucination (training-free):** Direct sequence optimization via backpropagation through a frozen struc-ture predictor, targeting high-confidence scores such as pLDDT or ipTM. Representative tools include BindCraft [38, 39] and BoltzDesign1 [13].

**Figure 1.**
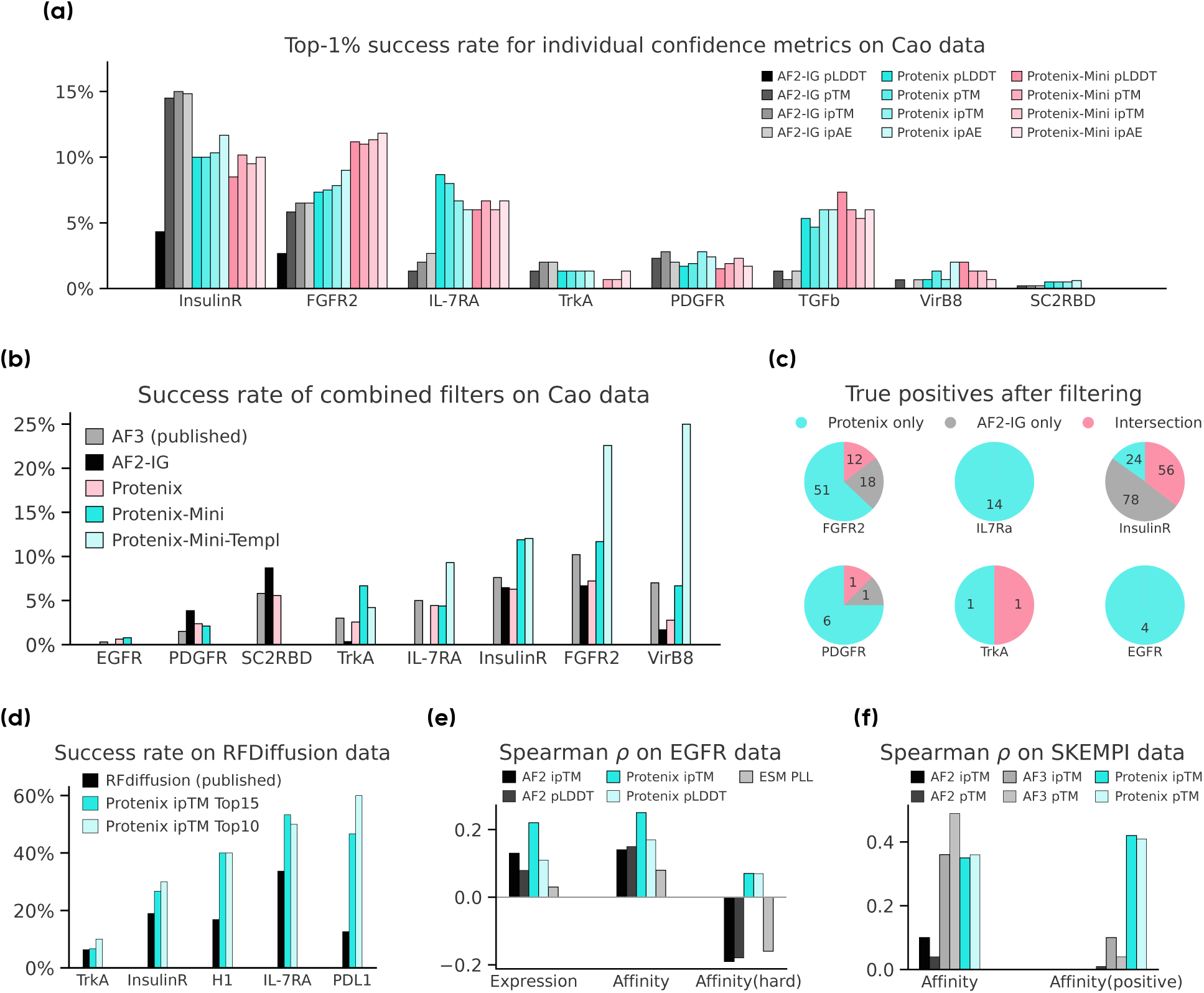
Filtering and ranking power comparison. **(a)** Top-1% success rates for individual confidence scores derived from AF2-IG, Protenix, and Protenix-Mini(-Templ) on Cao data. **(b)** Combined filter success rates across targets. “AF3 (published)” refers to results reported by Zambaldi et al. [59], while other results are computed via our unified pipeline. Filter thresholds for each model are listed in Table 2. **(c)** Venn diagrams show limited overlap between true positives retained by Protenix and AF2-IG filters, highlighting complementary design space coverage. **(d)** Re-ranking RFDiffusion binders [54] using Protenix ipTM improves success rates across all targets. RFDiffusion [54] generated 95 binder designs per target across five targets. “RFDiffusion (published)” shows the original experimental success rates, based on filtering with AF2-IG. “Protenix ipTM Top10(15)” reports success rates after re-ranking the same designs by Protenix ipTM and selecting the top 10 or 15, consistently improving hit rates across all targets. **(e)** On the EGFR competition dataset [16], Protenix better ranks expression and affinity, especially among binders with measurable affinity (“Affinity(hard)”). **(f)** On the subset of SKEMPI2.0 [25, 34], Protenix outperforms AF2 and matches AF3 [34] in ranking correlation. On positive-binding mutations (“Affinity(positive)”), Protenix shows pronounced advantage.

Both strategies have shown strong empirical results, yet few studies have systematically compared them under matched conditions. Here, we present a side-by-side evaluation of diffusion- and hallucination-based generation, with both methods implemented and optimized in-house to enable a fair comparison. In this section, we demonstrate that each method achieves state-of-the-art performance on *in silico* metrics, outperforming existing baselines. Building on this foundation, we compare the two strategies directly, highlighting their respective strengths and trade-offs for general-purpose protein binder design.

### 3.1 Development of Diffusion and Hallucination

**Diffusion.** We developed the protein design model, named PXDesign-d, built upon the Protenix structure prediction framework. PXDesign-d supports accurate, rapid, and programmable structure-based protein binder design. Details are described in Appendix C. Several key enhancements distinguish our approach:

- <u>Efficient architecture:</u> Unlike prior diffusion-based methods that rely on *SE*(3)-equivariant models or AlphaFold2-style frame representations [28, 54, 59], PXDesign-d directly generates Cartesian atom co-ordinates. Inspired by AlphaFold3, it employs a Diffusion Transformer (DiT) backbone without using expensive triangle updates during diffusion. This architectural simplification offers several-fold speedup in long-sequence generation while maintaining high structural fidelity, enabling scalable virtual screening and design of large proteins.
- <u>Unified multi-target training:</u> Although this work focuses on designing protein binders for protein targets, PXDesign-d is trained to support a wide range of molecular target types, including proteins, small molecules, nucleic acids (DNA/RNA), and post-translational modifications.
- <u>Controllable generation:</u> PXDesign-d supports a wide range of conditional inputs, including multiple sequence alignments (MSAs), structural priors from the target, target-specific hotspots, and user-defined preferences such as secondary structure or solvent-accessible surface area (SASA). While all results in this work were generated without applying user-defined preferences, these capabilities make PXDesign-d adaptable for future tasks requiring controllable generation.

**Hallucination.** We implemented PXDesign-h, a gradient-based sequence optimization pipeline using frozen Protenix predictors (Appendix B). Several enhancements are introduced to improve this approach:

- <u>End-to-end differentiation:</u> AlphaFold3-style models use a 200-step diffusion decoder by default, making backpropagation infeasible. We reduce this to a 2-step ODE-based sampler in Protenix [12], enabling efficient end-to-end differentiation through the entire structure prediction process.
- <u>Protenix-Mini for faster optimization:</u> We develop Protenix-Mini [23], a lightweight variant that offers similar predictive performance to the full model but significantly faster runtime, making it ideal for iterative sequence optimization.
- <u>Ensemble for robustness:</u> To improve generalization and avoid overfitting to any single predictor, we optimize sequences against an ensemble of five Protenix-based models, sampled randomly at each step.

### 3.2 In silico Benchmarks

We evaluate our model under two distinct settings: **unconditional protein monomer generation** and **conditional protein binder generation**. The unconditional generation task serves to demonstrate the model’s general capability to generate diverse and structurally realistic proteins without external constraints. In contrast, the conditional generation task reflects real-world applications in binder design, where the goal is to generate protein sequences and structures that can bind to a given target with high affinity and specificity.

#### 3.2.1 Unconditional Protein Monomer Benchmark

PXDesign-d outperforms or matches prior baselines (RFdiffusion [54], MultiFlow [10], Proteina [22]) in both designability and diversity across various lengths up to 1400 residues, with the performance gap widening as sequence length increases. Despite being trained only with a crop size of 640 residues, PXDesign-d maintains strong structural fidelity and diversity even for sequences exceeding 1000 residues, where competing methods show substantial degradation (Figure 2a). We note that recent works [17, 21], released after our benchmarking was completed, report improved performance on similar tasks. These results reflect the rapid progress in the field. Detailed datasets, evaluation metrics, and sampling hyperparameters are provided in Appendix D.

**Figure 2.**
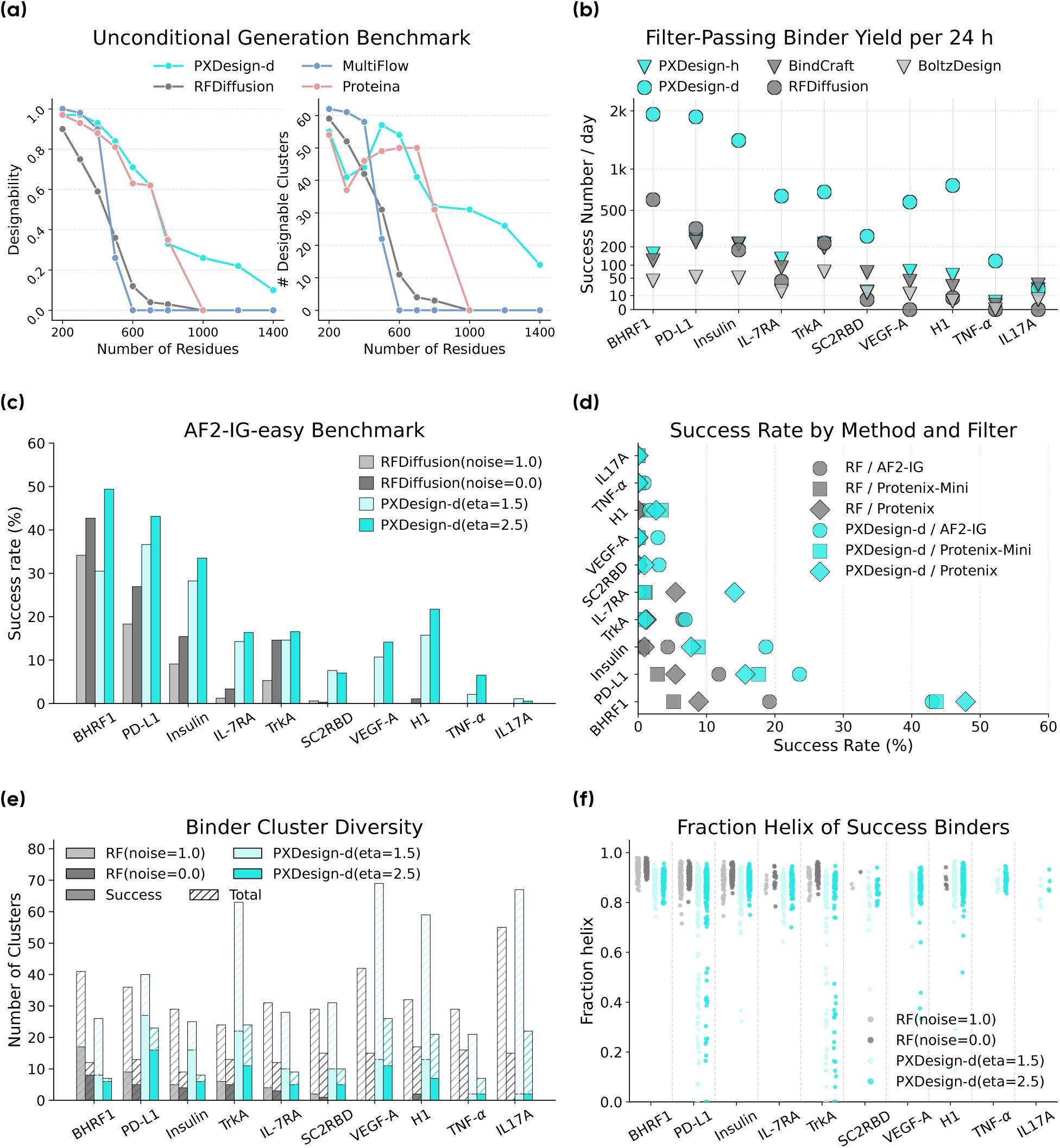
In silico evaluation of PXDesign on unconditional protein monomer design and conditional protein binder design tasks. **(a)** Unconditional monomer benchmark. Left: The ratio of designable proteins (scRMSD < 2). Right: The number of desigable structure clusters (TM-score < 0.5). **(b)** The time required for PXDesign-h and PXDesign-d to generate one sample passing AF2-IG-easy under default settings respectively. **(c)** Success rate of binders under the AF2-IG-easy filter defined in Table 2 across different methods on 10 representative protein targets. **(d)** Success rate of binders across different methods and filters defined in Table 2. **(e)** The number of structure clusters (TM-score < 0.5) of all binders and successful binders under AF2-IG-easy filter. **(f)** The fraction *α*-helix of successful binders.

#### 3.2.2 Conditional Protein Binder Benchmark

To extensively evaluate our method, following previous works [59], we use 10 protein targets with diverse structural properties as test set. These targets are not only biologically important, but also cover the difficulty of successfully designing binders for the proteins in the Protein Data Bank (PDB). Full benchmarking protocols, including dataset definitions, evaluation metrics, and runtime efficiency comparisons with hallucination-based methods, are described in Appendix D.

**Quality.** Across this benchmark, PXDesign-d consistently achieves higher success rates than RFdiffusion on the AF2-IG-easy criterion (Figure 2c). When applying alternative filters such as AF2-IG, Protenix-Mini, and Protenix (Figure 2d), PXDesign-d maintains both higher mean success rates and broader target coverage, with notable advantages on challenging cases like VEGF-A, IL17A, and TNF-*α*.

**Structural Diversity.** PXDesign-d generates a greater number of distinct structural clusters than RFdiffusion for nearly all targets (Figure 2e). Analysis of secondary structure composition (Figure 2f) reveals that PXDesign-d produces binders spanning a broad range of *α*-helix content, while RFdiffusion outputs are heavily *α*-helix-biased. This indicates broader coverage of fold space and greater structural versatility.

**Diffusion vs. Hallucination.** We directly compare our diffusion-based PXDesign-d with hallucination-based approaches, including our Protenix-powered PXDesign-h, BindCraft, and BoltzDesign1, under identical evaluation settings. Runtime analysis (Figure 2g) shows that PXDesign-d delivers more successful designs within 24h than any hallucination method, owing to its faster generation speed and higher pass rates. While hallucination remains competitive for targeted, small-scale optimization, diffusion is better suited for largez.

Given these advantages, all subsequent wet-lab experiments and the PXDesign webserver deployment are based on PXDesign-d (diffusion).

## 4 In vitro Experiments

We experimentally validated PXDesign on seven protein targets: Interleukin-7 Receptor Alpha (IL-7RA), SARS-CoV-2 receptor-binding domain (SC2RBD), Programmed Death-Ligand 1 (PD-L1), Tropomyosin receptor kinase A (TrkA), Vascular Endothelial Growth Factor A (VEGF-A), Epidermal Growth Factor Receptor (EGFR), and Tumor Necrosis Factor-*α* (TNF-*α*), chosen for both biological relevance and diverse design challenges (Table 3).

**Table 3.** Targets and experimental results of PXDesign-d binders. For SC2RBD and IL-7RA, “v0” indicates results from the initial PXDesign version. A binder is counted as successful if *K*_D_ *<* 1000 nM.

| Target | PDB ID | Crop | Hot-spot | Candidates Designed | Successful Binders |
| --- | --- | --- | --- | --- | --- |
| IL-7RA | 3DI3 | B17-209 | B58, B80, B139 | 10 (v0) | 4 (v0) |
| SC2RBD | 6M0J | E333-526 | E485, E489, E494, E500, E505 | 10 (v0) + 8 | 2 (v0) + 4 |
| PD-L1 | 5O45 | A17-132 | A56, A115, A123 | 17 | 14 |
| TrkA | 1WWW | X282-382 | X294, X296, X333 | 18 | 3 |
| VEGF-A | 1BJ1 | V14-107, W14-107 | W81, W83, W91 | 18 | 7 |
| TNF-α | 1TNF | A12-157, B12-157, C12-157 | A31, A32, A113, C73, C87 | 20 | 0 |
| EGFR | 1MOX | A1-186, A311-500 | A357, A380, A412 | 18 | 3 |

For each target, we used the PXDesign-d to generate a diverse set of *in silico* designs (60–160 aa) and applied a filtering pipeline: Candidates were required to pass both AF2-IG and Protenix filters (the Protenix ipTM cutoff was relaxed to 0.80 for VEGF-A, SC2RBD, and TNF-*α* to maintain diversity). Passing designs were clustered by structural similarity using Foldseek [49], and the highest-ranked representative by Protenix ipTM from each cluster was selected, yielding 8–18 candidates per target.

Selected binders were expressed in an *E. coli* cell-free system with an N-terminal Strep-tag and purified via Strep-Tactin affinity chromatography. Designed proteins with expression yields greater than 0.3 mg/mL (determined by A280) or 0.2 mg/mL (determined by the Bradford assay[8]) were subjected to BLI affinity measurements. An initial screening using biolayer interferometry (BLI) was conducted at a single concentration of 1000 nM to identify potential binders. Candidates exhibiting response signals above 0.06 nm were selected for multi-concentration BLI assays to determine *K*_D_. Binders with *K*_D_ *<* 1000 nM were considered successful. Both expression and BLI assays were performed by GenScript (Nanjing, China), and target proteins were purchased from Sino Biological (Beijing, China). For PD-L1 and EGFR binders, expression and affinity assays were performed at Sino Biological (Beijing, China). Due to differences in expression systems, the number of successfully expressed PD-L1 binders increased from 11 to 16. Additionally, the initial screening concentration was increased to 5 *µ*M to ensure the PD-L1 positive control met the screening criteria. Under this standard, the affinity of a PD-L1 mini-protein, which had been excluded during the previous single-concentration BLI, was successfully determined.

PXDesign can design structurally diverse protein binders that exhibit potent target binding (Figure 3), and achieve high-affinity binding. PXDesign v0 has already achieved strong performance on IL-7RA, surpassing all baselines except Chai-2. On SC2RBD, the updated PXDesign shows a marked improvement over v0, increasing the hit rate from 20.0% to 50.0%. Moreover, for PD-L1, PXDesign achieved a hit rate of 82.4% (14 out of 17), which is comparable to BindCraft (≈77.8%) and significantly outperforms all other baselines. Additionally, most of our binders exhibit *K*_D_ values below 100 nM; in contrast, the strongest binder designed by BindCraft has a *K*_D_ of approximately 1000 nM (Appendix E).

**Figure 3.**
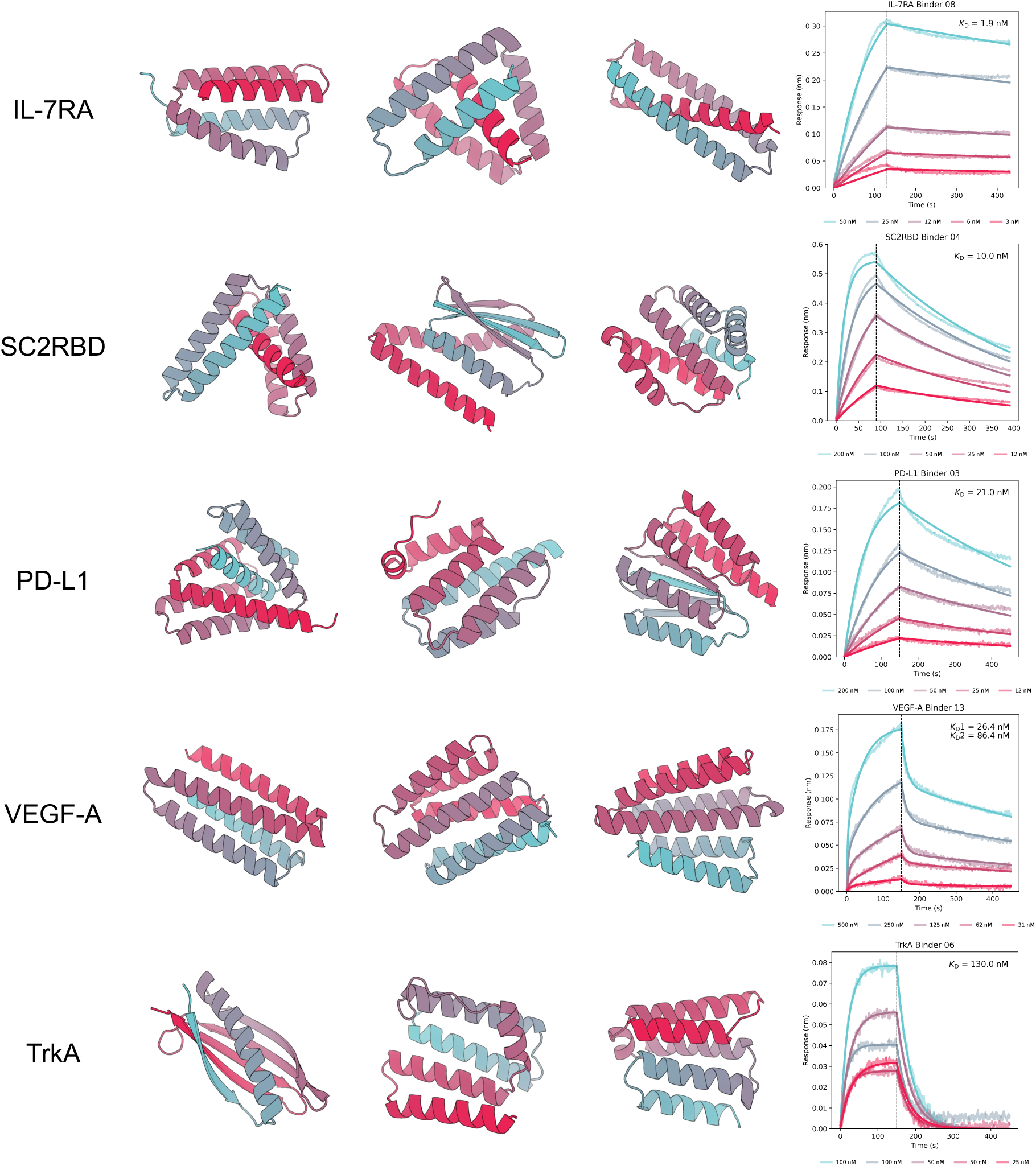
Designed binders and characterization. For each row, three structures of nanomolar binders were showcased. The right columns are BLI sensorgrams at multiple analyte concentrations; the vertical dashed line marks the switch from association to dissociation phases.

## 5 Limitations and Challenges

While confidence-based filtering shows strong potential, several challenges remain.

**Score variability across targets.** Confidence scores exhibit substantial variability across targets (Figure 4), which makes tuning and selecting unified thresholds challenging. We further observe that success rates among different targets have <u>Pareto tradeoffs</u>: improvements on one target often degrade performance on others, and Pareto improvements are difficult to achieve. For example, in Figure 4, raising the pTM threshold from 0.75 to 0.92 boosts the FGFR2 success rate from 5% to 100%, while the SC2RBD success rate falls from 0.1% to 0%. Effective filtering strategies must account for such target-specific shifts, as well as the constraints of limited evaluation datasets. Model-agreement-based approaches may help mitigate some of these weaknesses.

**Figure 4.**
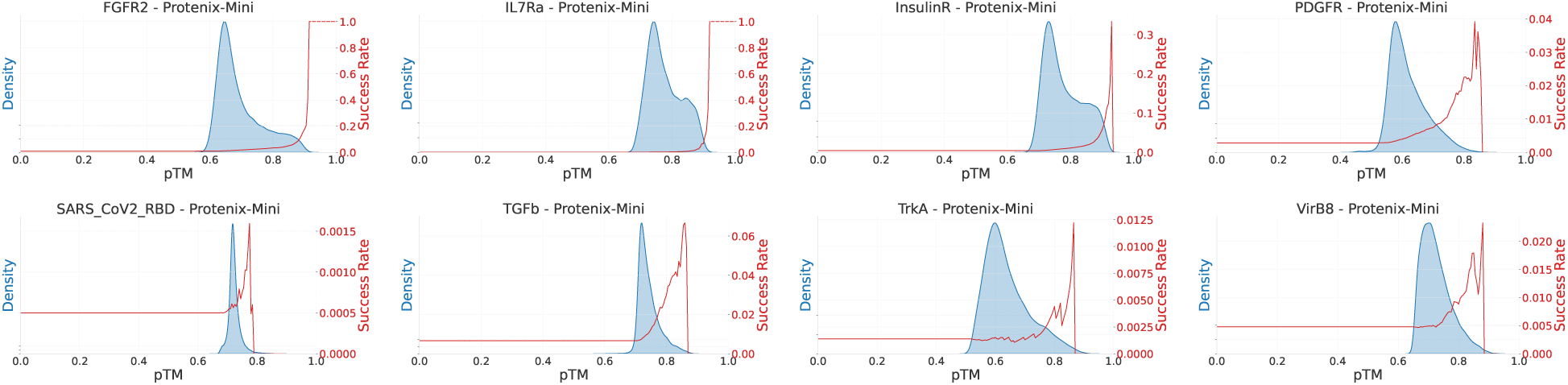
The Protenix-Mini pTM score distributions and success rates on different targets. The left y-axis shows the density of pTM scores, while the right y-axis displays the success rate across different pTM thresholds. Across different targets, the best success rates correspond to distinct pTM thresholds.

**Dataset limitations.** The Cao data contains very few strong binders, which limits its utility for evaluating high-precision filtering strategies. In particular, certain filters, such as Protenix-Mini on SC2RBD, exhibit zero success rate (SR) due to the sparsity of validated positives. While Protenix also shows low SR on the EGFR subset of Cao data (Figure 1b), it achieves substantially higher AUC and success rate on the EGFR competition dataset (Figure 9). Additionally, our wet-lab experiments indicate that intersection filters can be effective in practice, even though they appear overly stringent on Cao data. Despite these issues, Cao remains one of the only large-scale, publicly available benchmarks for binder filtering, underscoring the need for richer datasets to support fair and consistent evaluation.

**Pipeline limitations.** Our current pipeline relies on multiple sequential steps—generation, structural prediction, filtering, and clustering—each with its own computational cost and potential error propagation. While this modular design improves interpretability and flexibility, it can be resource-intensive for very large design spaces and may limit throughput in time-sensitive campaigns. Integrating filtering more tightly with generation or exploring early-stage, low-cost triage methods could improve scalability.

**Experimental limitations.** Our current experimental assessment of success rate is constrained by the throughput of the BLI assay. Incorporating display-based screening methods could substantially increase throughput, enabling the evaluation of approximately 100 designs per target. Similar to AlphaProteo, our pipeline encountered difficulty with one challenging target, TNF-*α*, despite achieving remarkable improvements in success rates for the other five targets. For PD-L1, six designs were excluded from testing due to low expression yields; however, nearly all of the remaining designs were confirmed to be functional binders. In addition to failures among our own designs, we occasionally encountered positive controls that did not successfully expressed. We are actively developing a robust experimental platform to enable reliable characterization of designed binders at scale.

## 6 From Protein Binders to a Unified Model for Molecular Design

While our current *in silico* and wet-lab validations focus on protein binder design, the underlying framework is readily extensible to a broad range of molecular targets. Systematic benchmarking and experimental validation of these additional modalities represent important future directions.

### 6.1 Demonstration Across Modalities

While the primary focus of this work has been on protein binder design, our design model naturally extends beyond these tasks. Since both our diffusion and hallucination generators are built on (or closely derived from) the Protenix structure prediction framework, they inherit the ability to model diverse biomolecular targets, including nucleic acids, small molecules, and post-translationally modified proteins. In principle, the same generative–filtering pipeline can be applied to these modalities.

At present, these broader applications have not undergone the same level of rigorous evaluation as protein binders. We have conducted case studies (Figure 5 and Figure 6c) that illustrate the feasibility of generating binders for nucleic acids, small molecules, and cyclic peptides, but these remain qualitative demonstrations.

**Figure 5.**
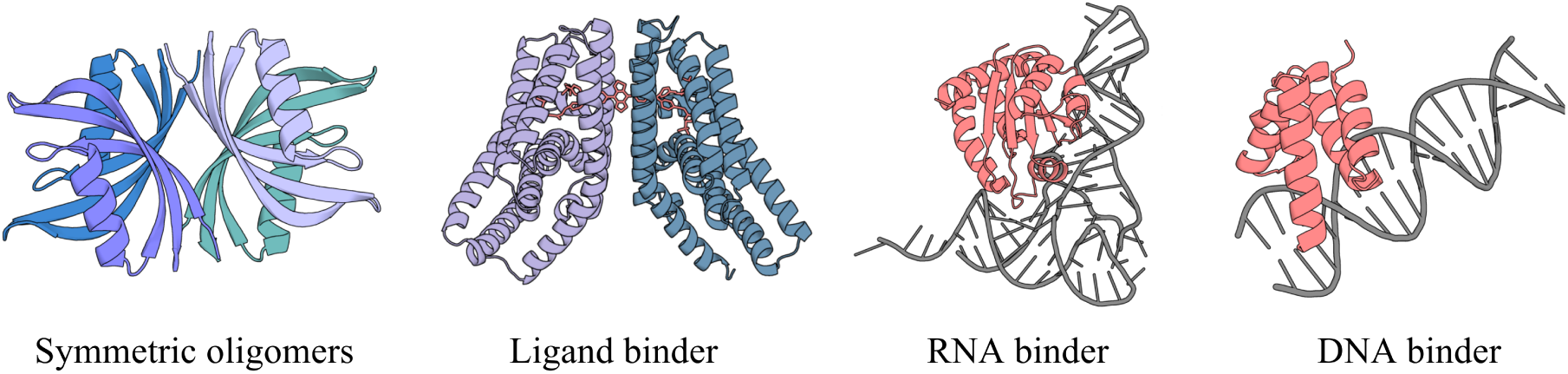
Case studies illustrating the versatility of our unified design model. Example designs for symmetric oligomers, ligand binder, RNA binder, DNA binder.

**Figure 6.**
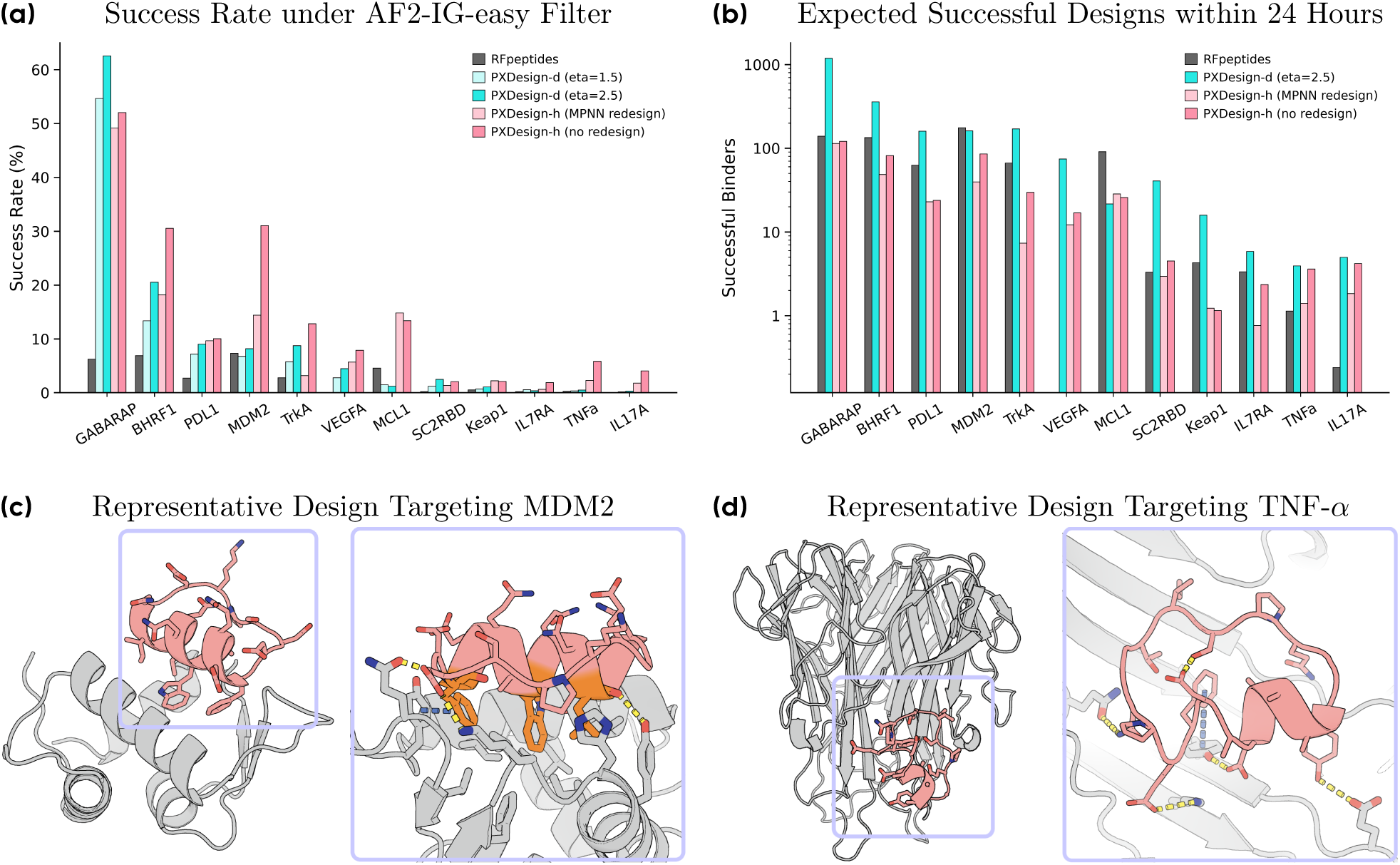
In silico evaluation of PXDesign on zero-shot cyclic peptide binder design. **(a)** Success rates of various methods across targets, evaluated using the AF2-IG-easy filter. **(b)** Expected number of successful cyclic peptide binders generated within a 24-hour period for each method and target. **(c)** Representative design targeting MDM2. Left: Protenix-predicted complex structure. Right: zoomed-in view of the binding interface. The conserved “F—W–L” triad is highlighted in orange. Key interactions are shown. **(d)** Representative design targeting TNF-*α*. Left: Protenix-predicted complex structure. Right: zoomed-in view of the binding interface. Key interactions are shown.

### 6.2 Cyclic Peptide Binder Benchmark

Cyclic peptides are emerging as a promising therapeutic modality due to their enhanced membrane permeability, oral bioavailability, synthetic tractability, low immunogenicity, and capacity for specific binding to protein surfaces previously considered “undruggable” [26, 29, 37, 41, 41, 42, 42, 51–53, 60]. Cyclic peptide binder design is structurally and biophysically similar to protein binder design, allowing reuse of *in silico* metrics with minimal modification. As a first cross-modality benchmark, we extended both PXDesign-d and PXDesign-h for zero-shot cyclic peptide binder generation targeting 12 diverse proteins, including AlphaProteo targets [59] and proteins with previously reported binders [41, 42], spanning sequence lengths of 8–18 residues.

Our models consistently outperform RFpeptides [41] in success rates under the AF2-IG-easy criterion (Figure 6a) and in efficiency measured by the expected number of successful designs per 24h (Figure 6b), with particularly strong gains on challenging targets such as TNF-*α*. While PXDesign-h is slower, it achieves superior performance on specific targets (e.g., MDM2, MCL1, IL17A, TNF-*α*), and its performance further improves when using native sequences without ProteinMPNN redesign—highlighting its ability to generate viable sequences directly (Figure 12).

Detailed benchmarking protocols and per-length analyses are given in Appendix F. Representative designs for MDM2 and TNF-*α* (Figure 6c–d) illustrate structurally plausible interfaces, with the MDM2 design recovering the conserved “F—W–L” interaction triad critical for hydrophobic binding [7].

## 7 Discussion

We have presented PXDesign, a unified framework for structure-based molecular design that integrates scalable diffusion-based generation (PXDesign-d) with complementary hallucination-based search, coupled to multi-filter prioritization using orthogonal structure predictors (AF2-IG and Protenix). Across extensive *in silico* benchmarks and wet-lab validation on seven biologically diverse protein targets, PXDesign-d demonstrates higher throughput, higher pass rates, and broader structural diversity than hallucination approaches, making it well-suited for large-scale exploratory campaigns. Guided by these findings, both our experimental pipeline and public web server are currently built on PXDesign-d.

Our filtering analyses reveal that no single confidence metric generalizes across all targets—success rate optima vary, and ensemble filtering provides broader design space coverage. Current public benchmarks, such as the Cao dataset, have sparse positive signal, constraining statistical resolution and sometimes misrepresenting practical performance. Improved, community-shared datasets with richer annotations are essential for accelerating method development and enabling fair comparisons.

While this work has focused on protein–protein binder design, the framework generalizes naturally to other modalities. Our case studies and initial benchmarks on cyclic peptide binders demonstrate that minimal modifications allow zero-shot generalization beyond proteins, and the same generative–filtering pipeline is in principle applicable to nucleic acids, small molecules, and post-translationally modified proteins. Scaling experimental validation to these modalities—particularly where high-quality benchmarks and structure predictors are available—is an important next step.

From a modeling standpoint, our results underscore the advantages of generative architectures that are both high-throughput and conditioning-flexible. Diffusion provides efficient large-scale sampling, while hallucination retains value for targeted, small-scale optimization. As structure prediction continues to advance, deeper integration of prediction and design—where predictors act not only as filters but also as gradient-informed guides—could enable faster, more accurate, and more interpretable workflows. Protenix already illustrates this potential by serving both as a scoring function and as a backbone for generative models.

In summary, PXDesign offers a practical and extensible pipeline for large-scale structure-based design, validated in wet-lab experiments and accessible via a public web server. By combining complementary generative paradigms, orthogonal filtering, and open benchmarks, we aim to support a broader shift toward unified, general-purpose molecular design systems that bridge computational discovery and experimental realization.

## 8 Revision History

### Version 3 (December 12, 2025)

- Updated In Vitro Results:

**–** Added EGFR as an additional validation target.

**–** Updated PD-L1 results following independent validation by a CRO. The revised experiments resolved previous yield issues, enabling successful expression of most designed binders.

**–** Adopted an end-to-end hit rate calculation for results of PXDesign, defining the hit rate as the fraction of binders with *K*_D_ *<* 1000 nM among all experimentally tested designs (including those not expressed) to provide a strict assessment of the generative pipeline’s real-world success.

**–** Added Boltzgen as an additional comparison method.

**–** Included exemplary *K*_D_ values from our measurements for both positive controls and PXDesign-generated binders.

**–** Updated the experimental hit rates for AlphaProteo and RFdiffusion to be fully consistent with the values reported in the respective original publications. For Latent-X, we additionally report hit rates based on the number of binders with measured *K*_D_ values within 1000 nM. To improve readability, hit-rate percentages are reported as integers.

- Fixed minor typos and updated several citations.

### Version 2 (September 2, 2025)

- Added missing citations.
- Added details for benchmarking configurations.
- Updated BLI sensorgrams in Figure 3 with better fit.
- Added the “Protenix-basic” filter criteria in Table 2, which is used for some challenging targets in our wet-lab experiments.

### Version 1 (August 16, 2025)

Initial maniscript.

## Acknowledgements

We thank members of our team for their support, including Wenbo Lin, Jiazheng Zhou, and Fangzhou Guo for their work on webserver development. This project builds upon the internal Protenix codebase^1^, and we thank all contributors to its development.

## Footnotes

1 https://github.com/bytedance/Protenix

## Appendix

### **A** Filtering Methodology and Benchmark Evaluation

#### Filter Development

To identify general-purpose, model-specific filters, we perform a grid search over combinations of confidence scores. We construct and evaluate confidence-based filtering strategies from two structure predictors: **Protenix** [12] and **AF2-IG** [54]. AF2-IG is a variant of AlphaFold2 adapted for binder design where “IG” denotes “initial guess” [5, 54]. We also include **Protenix-Mini**, a compact variant with reduced model size, and **Protenix-Mini-Templ**, which uses the target structure as a fixed template without relying on MSA input. All Protenix models utilize a 2-step ODE sampler for efficiency, replacing the standard 200-step diffusion sampler.

We design a two-stage filtering strategy to identify high-quality protein complex predictions generated by Protenix. In the first stage, we identify informative evaluation metrics and select an optimal triplet combination based on their ranking performance across targets. In the second stage, we perform a grid search over the cutoff values of the selected metrics to determine optimal thresholds for filtering.

**Stage 1: Metric Selection and Combination.** We first evaluate the ranking performance of individual metrics using a Top-1% threshold classifier and compute the Enrichment Factor (EF) to measure how effectively each metric identifies high-quality predictions. For each target, we select the Top-3 metrics with the highest EF values. Based on frequency analysis, we identify eight most frequently selected metrics (some with the same EF score): binder pTM, complex pLDDT, interface pLDDT, binder ipTM, complex pTM, binder chain pLDDT, complex gpDE, and interface gpDE. From these eight metrics, we evaluate all possible triplets (*C*(8, 3) = 56 combinations) as filters. For each triplet, we apply the Top-1% quantile as a threshold and classify a sample as positive only if it satisfies all three thresholds simultaneously. We compute the Success Rate (SR) for each combination on every target and rank them accordingly. The final optimal combination is determined by aggregating rankings across all targets and selecting the one that achieves the most “Top-1” positions. The final optimal combination is determined by aggregating rankings across all targets and selecting the one that achieves the highest ranking among all targets. Ultimately, we find that a simplified filter using only two metrics, binder pTM and binder ipTM, achieves comparable or better performance than the full triplet, while offering improved robustness and interpretability. Therefore, we adopt {binder pTM, binder ipTM} as the final filtering metric set.

**Stage 2: Cutoff Grid Search.** Finally, to refine the filtering process, we perform a grid search to determine the optimal cutoff values for the two selected metrics, binder pTM and interface ipTM. We formulate the filter selection problem as an optimization problem, max (SR_1_(**x**), SR_2_(**x**)*, . . . ,* SR*_k_*(**x**)) ,

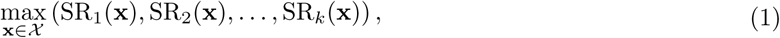

in which SR*_i_* denotes the success rate on the target *i*, and the set X is the feasible set of confidence score threshold combination. Typically, there is no feasible solution that can maximize all objective functions simultaneously. Consequently, the focus is the solutions where improving any objective cannot be achieved without deteriorating at least one other objective, which is defined as <u>Pareto Frontier</u>.

<u>Definition:</u> A solution *x*_1_ ∈ X dominates *x*_2_, if

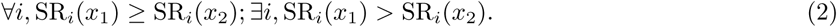

A solution *x*^∗^ ∈ X is Pareto optimal if there does not exist another solution *x* that dominates it. The set of Pareto optimal is called Pareto Frontier.

For each threshold combination, we compute the success rate (SR) on each target. Following the definition of Pareto Frontier, the search algorithm can come to a set of optimal points (as demonstrated in Figure 7). To distinguish the final solution, we tend to the solution which has minimal shifts across different targets, known as robust selection or risk aversion policy [4]. We calculate the rank of each combination’s SR within each target and then take the average rank across all targets. This average rank serves as the overall “score” for the threshold combination. As demonstrated in Figure 7c, we come to a balanced SR solution on FGFR2 and SC2RBD. Ultimately, we select the combination with the highest score as the final filtering criteria.

**Figure 7.**
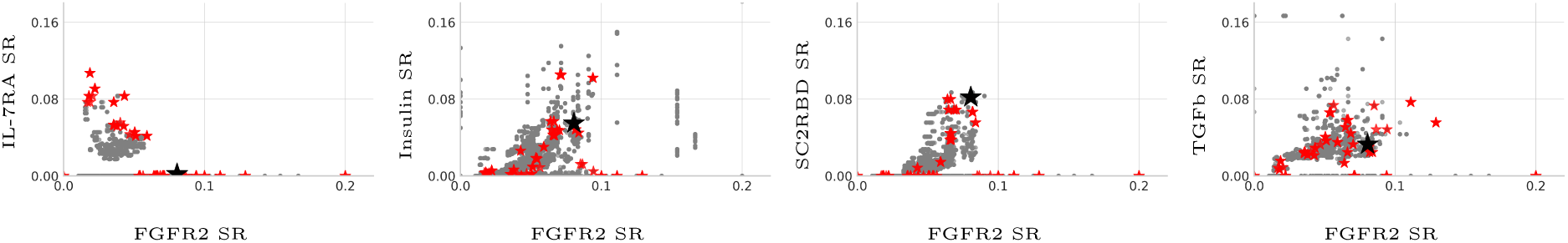
The performance of the AF2 confidence score filters. The SR for each confidence combination is plotted as one gray dot. The pareto frontier filters are highlighted as red stars, and the selected one is marked as a black star.

### Additional Benchmark Results

#### Per-Score Filter Evaluation

To complement the Top-1% SR analysis in the main text (Figure 8), we provide additional evaluation of individual confidence scores using two standard ranking metrics: AUC (area under the ROC curve) and AP (average precision, or area under the PR curve). These metrics reflect how well each score discriminates binders from non-binders across a range of thresholds, independent of any fixed selection cutoff.

**Figure 8.**
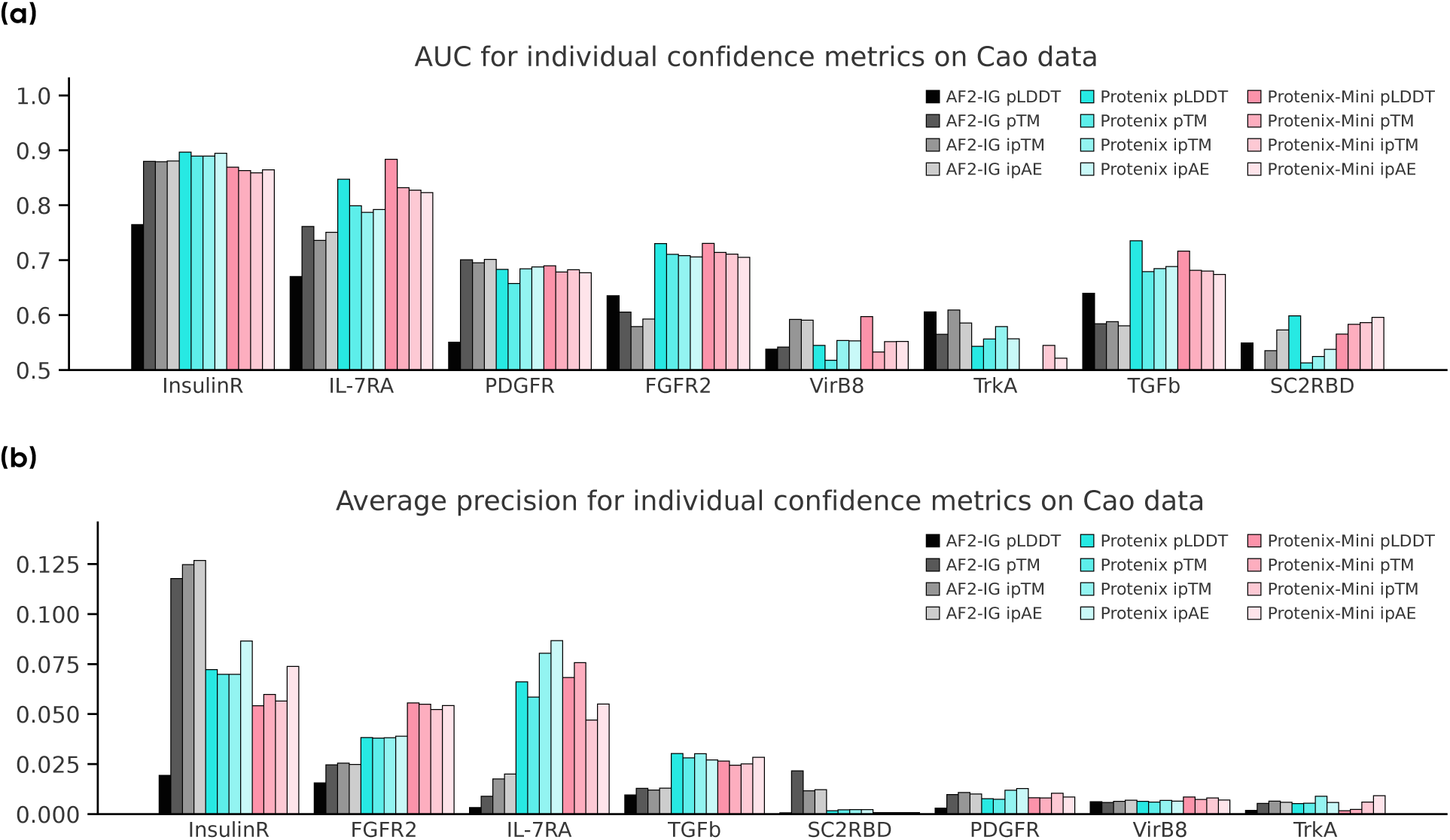
AUC and Average Precision scores for individual confidence metrics on Cao data. **(a)** Higher values indicate better global discrimination between binders and non-binders. **(b)** Similar to AUC, but more sensitive to top-ranking false positives.

We report AUC and average precision scores for individual confidence metrics across diverse design targets (Figure 8). These results are generally consistent with the Top-1% SR trends, reinforcing that Protenix-derived scores outperform AF2-based scores in most cases. However, no single metric is universally opti-mal—performance varies by target and model. For instance, AF2 shows stronger precision on Insulin and SC2RBD, as well as higher AUC on TrkA.

#### Ranking Accuracy

We further assess whether confidence scores can effectively prioritize designs by binding strength, beyond binary filtering. As shown in Figure 9, Protenix-derived scores consistently achieve higher AUCs across two distinct datasets: **(a)** On the EGFR binder challenge [16], which includes 400 designed binders with expression and binding affinity annotations, Protenix outperforms AF2 and ESM across multiple ranking metrics. **(b)** On the SKEMPIv2 subset filtered by AF3 [34], Protenix matches or exceeds AF3 in AUC overall and shows particularly strong performance on affinity-increasing mutations.

#### **B** PXDesign-h (hallucination) Details

Protenix, as a structure prediction model, not only excels at predicting complex protein structures but also provides reliable confidence estimates for those predictions. By treating the input binder sequence logits as trainable parameters, we can leverage these confidence scores to perform backpropagation through Protenix, effectively navigating the sequence space to discover high-quality binder candidates.

The hallucination process consists of two key components: (1) backbone models that supply gradient signals for updating the sequence parameters, and (2) optimization strategies that effectively utilize this gradient information. In this section, we first describe the backbone models employed in our framework, followed by an overview of the loss functions and optimization strategies used to guide the discovery of promising binder sequences.

#### Backbone structure prediction models

We employ five variants of the Protenix model family to provide gradient signals: Protenix, Protenix-Mini, Protenix-Mini-v2, Protenix-All-Data, and Protenix-Template. Among these, Protenix-Mini-v2 denotes a version of Protenix-Mini trained with different random seeds for its confidence module; Protenix-All-Data refers to a Protenix-Mini variant trained on a larger and more diverse dataset; and Protenix-Template indicates a Protenix-Mini model trained with the additional guidance of provided template structures. To prevent overfitting to any single model during the hallucination process, we randomly select one of these structure prediction models at each optimization step to perform the forward and backward passes, thereby generating the necessary gradient information for sequence updates.

#### Loss and Optimization

Thanks to its two-step diffusion architecture, the Protenix-based hallucinator enables end-to-end backpropagation of gradients from all confidence metrics, rather than being limited to contact loss derived from Pairformer outputs [13]. In our implementation, we combine multiple loss components with the following weights: pLDDT (0.15), pAE (0.4), ipAE (0.1), contact loss (1.0), interface contact loss (1.0), helix loss (−0.3), and radius of gyration of the designed binder (0.3). The core ingredients in hallucination process can be described as:

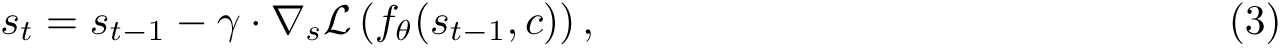

where *s_t_*represents binder sequence logits at time step *t*, L is the loss functions defined on the outputs of structure prediction model *f_θ_*, *c* denotes the conditions, e.g. target structures and sequences and *γ* is the step size for optimization. For binder hallucination, we adopt a 4-stage relaxed-to-hard sequence optimization process inspired by Pacesa et al. [38] and Cho et al. [13], which is detailed below.

*Stage 0: Softmax Warm-up* To avoid unstable exploration in the continuous logits space and to encourage the generation of realistic binder sequences, we begin with randomly initialized binder sequence logits. During this stage, we perform gradient descent updates guided by the backbone model’s feedback, followed by a softmax operation on the logits. This procedure ensures that the logits represent a valid amino acid distribution at each step.

*Stage 1: Soft logits update* We use logits from Stage 0 to re-initialize our binder sequence logits and start soft-logit updates during this stage. As the optimization steps increase, we continuously transfer binder sequence representation from soft logits *s* to softmax logits *p* = softmax(*s*) by linear interpolating soft logits and softmax logits. For example, we have in total *T* steps in this stage, at time step *t*, binder sequence is represented as (1 − *λ*)*s* + *λp*, where *λ* = *<u>^t^</u>*<u>^+1^</u> .

*Stage 2: Softmax logits update* In this stage, the goal is to transfer sequence logits representation from continuous representation to the final one-hot representation. To do so, we slowly decrease the temperature *τ* to get temperature-conditioned softmax logits *p* = softmax( *<u>^s^</u>* ). For a *T* -step stage 2, temperature at time step *t* is given by *τ_t_* = 0.01 + (1 − 0.01) × (1 − (*t* + 1)*/T* )^2^. Furthermore, we also use *τ_t_* to scale our learning rate *γ_t_* = *γ* · *τ_t_*to stabilize the final update of the binder sequence representation that are close to one-hot representation.

*Stage 3: One-hot sequence update* Finally, we directly get one-hot sequences by taking argmax from softmax sequence logits. To update one-hot sequence, we update softmax sequence using the gradients on it. The hard sequence *s^h^* at time step *t* is given by *s^h^* = stop_grad(argmax(softmax(*s_t_*)) − softmax(*s_t_*)) + softmax(*s_t_*).

### **C** PXDesign-d (diffusion) Details

#### Model Architecture

PXDesign-d is a diffusion-based protein design model built as a direct extension of the Protenix all-atom structure prediction framework. To enable generative capabilities, we introduce a special token [xpb] to denote residues to be designed. Each [xpb] token consists of four backbone atoms (N, CA, C, O). During training, the coordinates of all atoms are perturbed with noise and subsequently denoised through a learned diffusion process. Target residues are soft-conditioned through pairwise features. Specifically, the single-token condition *s* is initialized by embedding basic residue-level features, such as amino acid identity, hotspot annotations, etc. The pairwise condition *z* is initialized by embedding binned pairwise distances derived from the target structure. If no distance information is available for a residue pair, we assign it to a special bin.

Because binned pairwise distances offer strong structural constraints, we find it unnecessary to freeze the coordinates of target residues during training. Instead, the model learns to recover structure directly from these embedded pairwise signals. For the condition-based generation task, unlike previous methods [54] that relied on inpainting, PXDesign-d directly generates the coordinates of all atoms from noisy structure. Throughout the process, no additional constraints are imposed on the noise. This approach preserves the topology of the condition region while allowing flexibility in its side chains.

The overall architecture consists of two components: the prior module and the diffusion module. The diffusion network component includes 4 layers of atom-level attention encoders, 16 layers of a token-wise transformer, and 4 layers of atom-level attention decoders.

#### Training

PXDesign-d is trained on a large and diverse dataset that integrates both experimentally resolved structures and distilled data derived from predictive models. The dataset composition closely follows the training corpus used for Protenix, but with several extensions. We curated a subset of the Protein Data Bank (PDB) [6] up to May 1, 2021, and categorized complexes based on their molecular context. These include protein monomers as well as complexes involving proteins, ligands, DNA, or RNA. For each complex, we define task-specific design objectives by specifying which residues are subject to design, allowing for fine-grained control over sampling across different structural and functional categories. To enhance coverage and diversity, we supplement the experimental data with high-confidence structures predicted by AlphaFold2. Specifically, we incorporate monomer structures from the AlphaFold Protein Structure Database (AFDB) [50] and MGnify [43].

PXDesign-d is trained from scratch in two stages to facilitate both general structural modeling and target-conditioned generation. In the first stage, we upweight monomer-only distillation data, enabling the model to learn the fundamentals of protein backbone geometry in an unconditional setting. In the second stage, we gradually shift the sampling distribution toward experimentally resolved PDB complexes and target-conditioned design tasks.

The model is trained end-to-end by minimizing a weighted combination of loss functions. Specifically, we apply an MSE loss over all heavy-atom coordinates and a smooth LDDT loss, as used in Protenix for structure prediction. Additionally, we find that introducing a distogram loss on projected token embeddings helps improve local consistency and geometric plausibility. For the distogram loss and the smooth LDDT loss function, during training, we masked the samples with a noise scale less than 4Å. The total loss is:

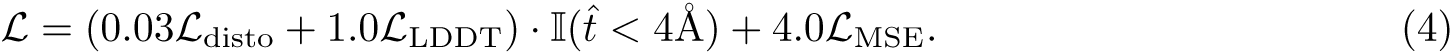

We use a crop size of 640 residues, a batch size of 64, and a diffusion batch size of 8. The model is optimized using Adam with a learning rate of 0.0005. The diffusion noise schedule follows the same formulation as Protenix.

#### Sampling and Evaluation in Benchmark

We perform diffusion-based sampling with 1000 steps for monomer generation and 400 steps for binder generation. We set the sampling parameters *γ*_0_ = 1.0 and *γ*_min_ = 0.01 via a preliminary hyperparameter grid search.

As shown in Figure 11b, the parameter *η* controls the trade-off between designability and diversity: higher values of *η* generally lead to higher-quality structures but reduced structural diversity. An ablation study of different *η* schedules shows that both linear and piecewise schedules outperform fixed-*η* variants, particularly for long-sequence generation. Based on these results, we adopt a piecewise schedule in our final model, where *η* = 1.0 for *t <* 0.65 and *η* = 2.0 for *t* ≥ 0.65.

**Figure 9.**
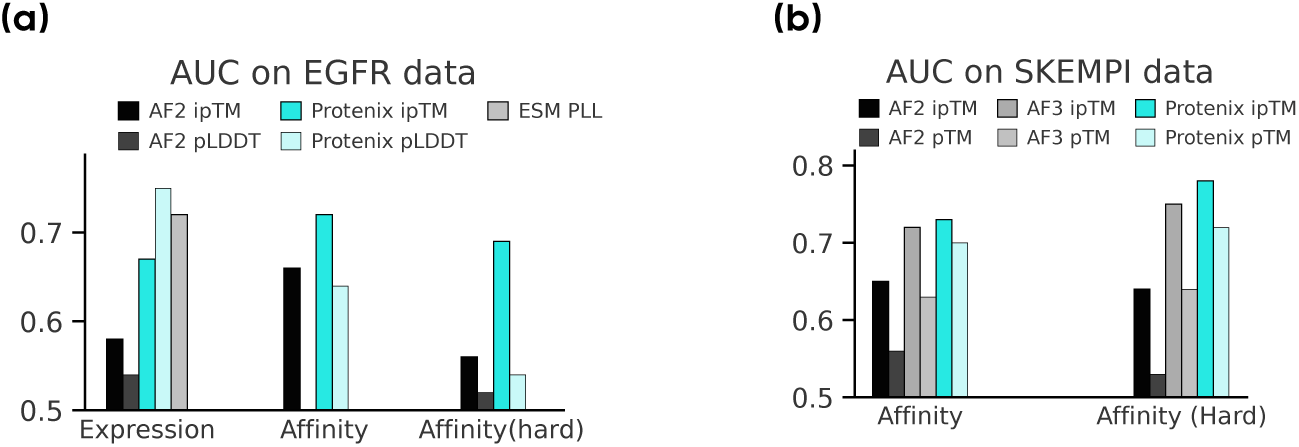
**AUC on EFGR and SKEMPI data.**

**Figure 10.**
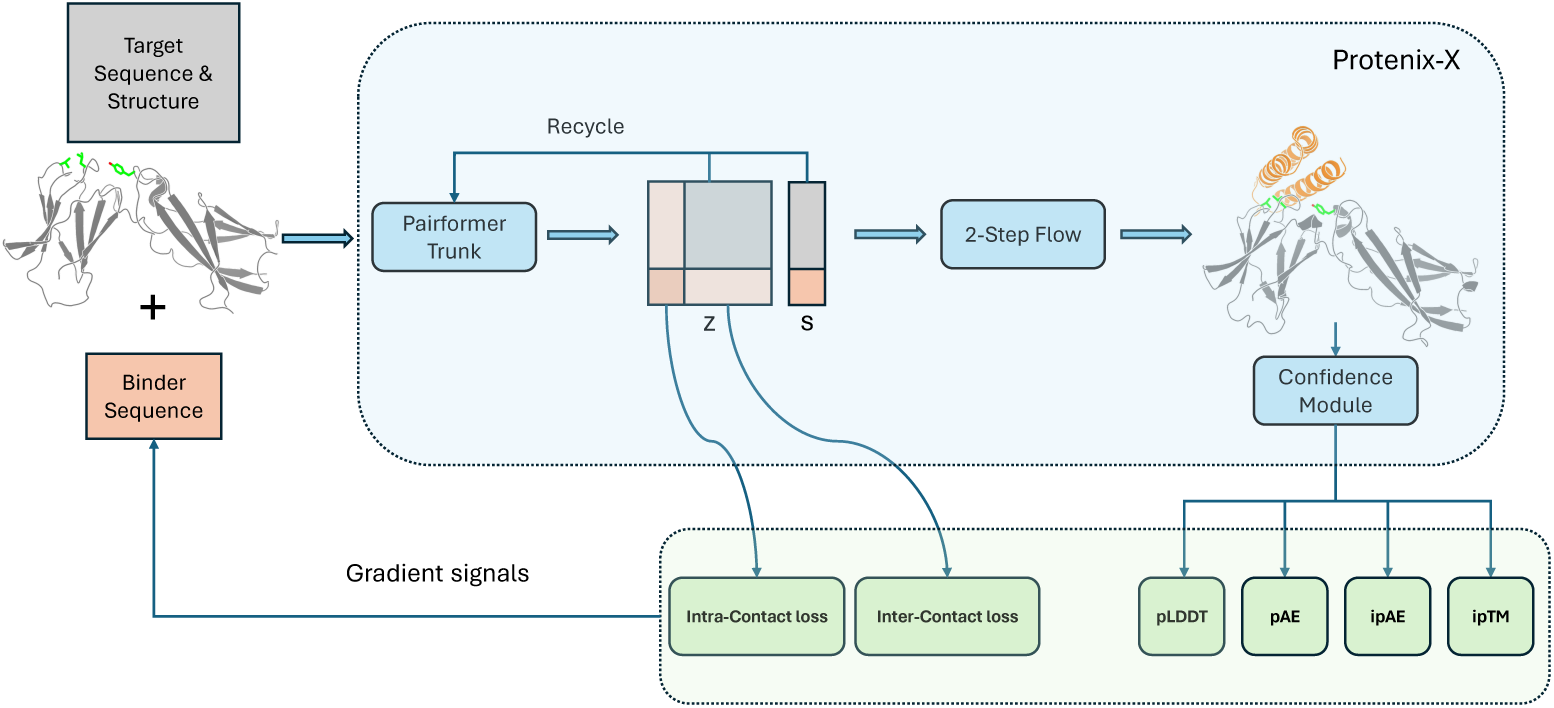
Overview of one-step optimization in Protenix-Hallucination. At each optimization step, a backbone structure prediction model, denoted as <u>Protenix-X</u>, is randomly selected from five candidates: <u>Protenix</u>, <u>Protenix-Mini</u>, <u>Protenix-Mini-v2</u>, <u>Protenix-Mini-All-Data</u>, and <u>Protenix-Mini-Template</u>, to prevent overfitting to the gradient pref-erences of any single model. During the forward pass, the target structure, target sequence, and binder sequence are input into <u>Protenix-X</u> to generate sequence representations *s* and pairwise representations *z*. Subsequently, an ODE-based sampler with 2 sampling steps efficiently predicts complex structures, and a loss defined on the generated structures is computed. Finally, gradients from this end-to-end differentiable process are used to update the protein binder sequence logits.

**Figure 11.**
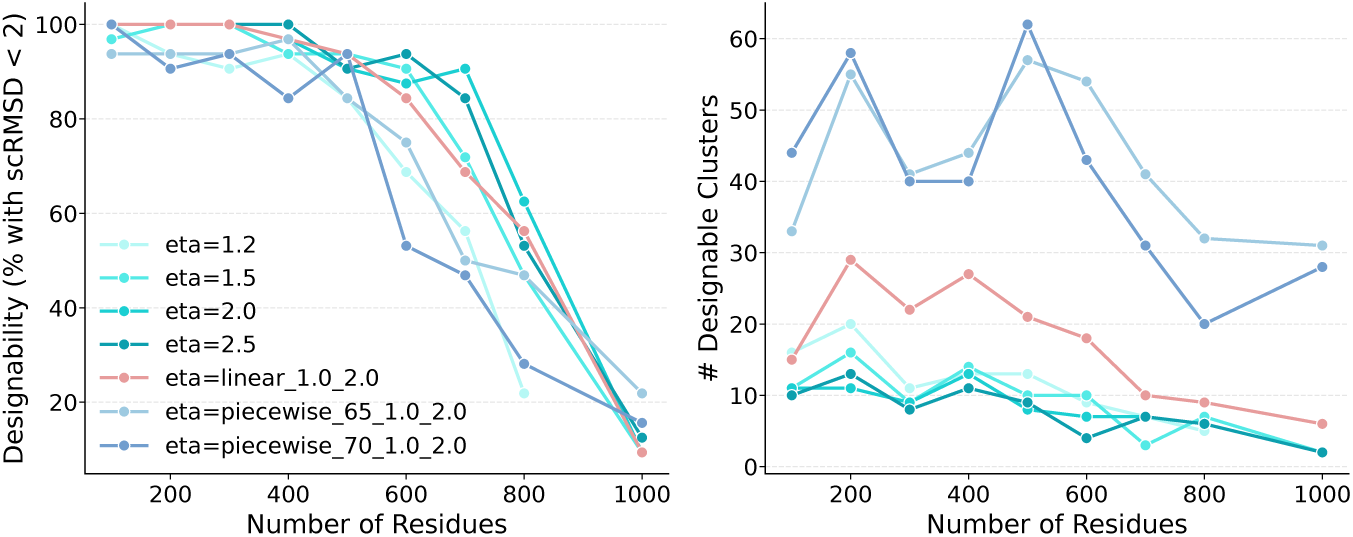
Ablation on *η* schedule in unconditional design. *η* increase from 1.0 to 2.0 in the linear and piecewise schedules. Piecewise schedules increase *η* at *t* = 0.65 or *t* = 0.7.

**Figure 12.**
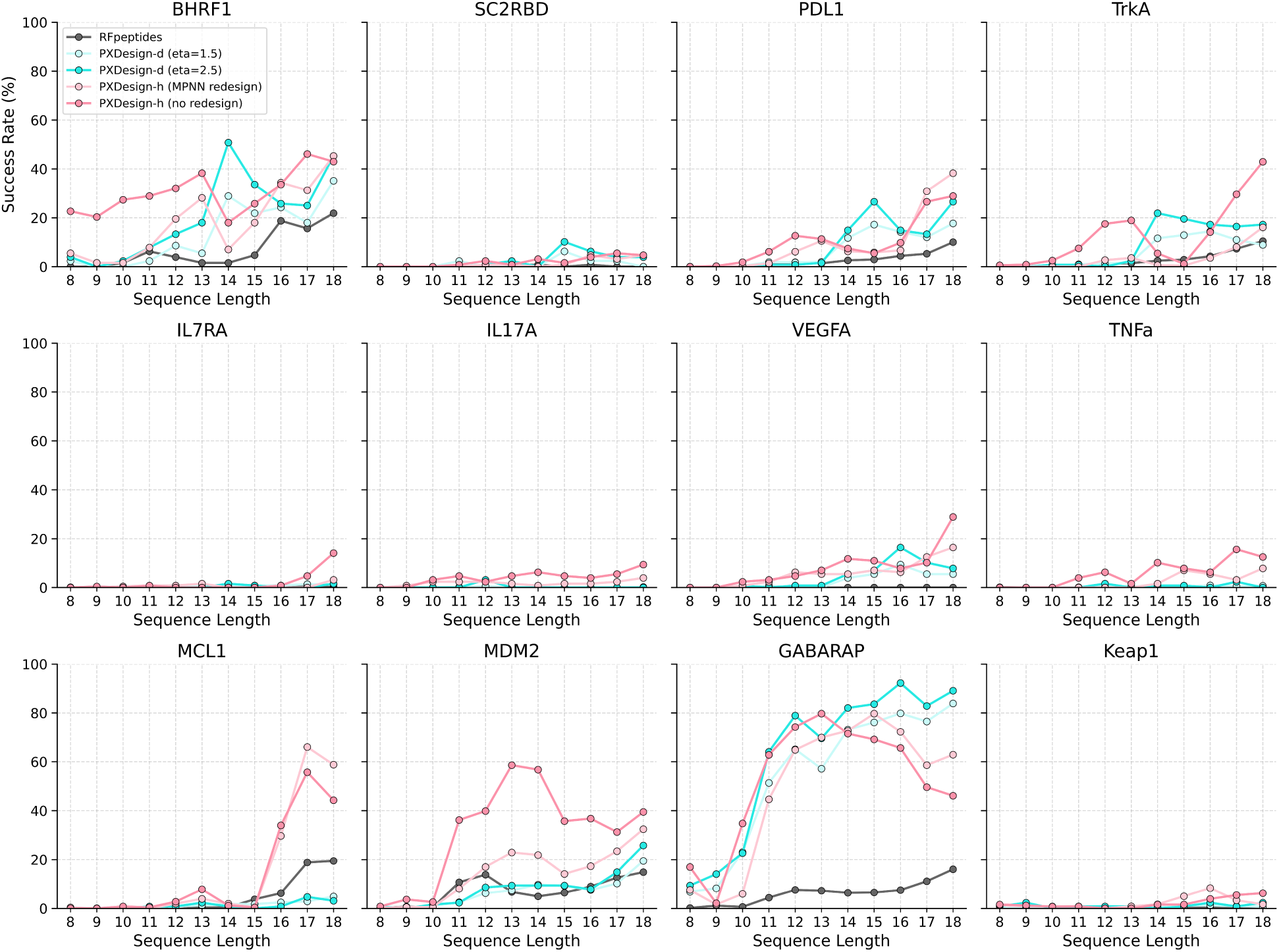
Per-length in silico success rates for zero-shot cyclic peptide binder design. Success rates are assessed under the AF2-IG-easy filtering criterion.

### **D** Generator Benchmarking Details

#### Unconditional Monomer Generation Protocol

For each sequence length (200–1400 residues), we generate 100 monomer backbones per method (RFdiffusion, MultiFlow, Proteina, PXDesign-d). Sequence design is performed using ProteinMPNN-CA [18] with default settings, followed by structure prediction using ESMFold [31]. Since PXDesign-h is specialized for binder design, we exclude it from this comparison.

**Designability metric**: A backbone is considered designable if the best self-consistency RMSD (scRMSD), computed over 8 independently designed sequences, is below 2 Å.

**Diversity metric**: Following Yim et al. [58], designable backbones are clustered using a TM-score threshold of 0.5, and the number of resulting clusters is reported.

The same evaluation protocol and random seeds are used across all methods to ensure fair comparison.

#### Conditional Binder Generation Protocol

Following Zambaldi et al. [59], we use a benchmark set of 10 protein targets with diverse structural properties. For each target, we generate binders using RFdiffusion (noise = 0.0 and 1.0), PXDesign-d, or hallucination-based methods, then perform sequence design with ProteinMPNN [18] at temperature 0.0001.

Since structure generation speed differs substantially between diffusion- and hallucination-based methods, we adopted evaluation settings that reflect realistic computational trade-offs. For diffusion-based designs, each generated structure was paired with a single ProteinMPNN sequence. For hallucination-based designs, where structure generation is much slower, we followed the BoltzDesign1 protocol [13] and sampled 8 sequences per structure to increase the chance of success. A design was deemed successful if at least one sequence passed the filter criteria defined in Table 2.

**Diversity metrics**: We cluster generated binders using TM-score < 0.5 and report both the number of clusters and the number of successful clusters.

**Secondary structure analysis**: Secondary structure content (*α*-helix percentage) is computed using DSSP on the folded structures, aggregated per target.

**Runtime efficiency**: To better reflect real-world efficiency, we measured both generation and evaluation times for different methods. Specifically, we measure the number of successful designs generated within 24 hours (including generation + evaluation) for diffusion- and hallucination-based methods (PXDesign-h, BindCraft, BoltzDesign1). we used the default experimental settings for each method. The length of generated binders strictly followed the previous work. For various targets, we generated 1,280 to 2,880 samples using NVIDIA A800 GPUs and recorded the average number of AF2-IG-easy filter-passing candidates produced within 24 hours.

**BindCraft adjustments**: Since BindCraft integrates hallucination and evaluation in a single pipeline, we removed evaluation time from our measurement to enable a fairer comparison. Specifically, we measured the time consumed by the binder_hallucination function (https://github.com/martinpacesa/BindCraft/blob/ main/bindcraft.py, Lines 109-111). Within this function, we counted only the time taken to hallucinate the binder, excluding the time spent on trajectory checking (https://github.com/martinpacesa/BindCraft/ blob/main/functions/colabdesign_utils.py, Lines 177-233).

A complete list of the GitHub repositories and commit hashes for all compared methods is provided in Table 4.

**Table 4.**
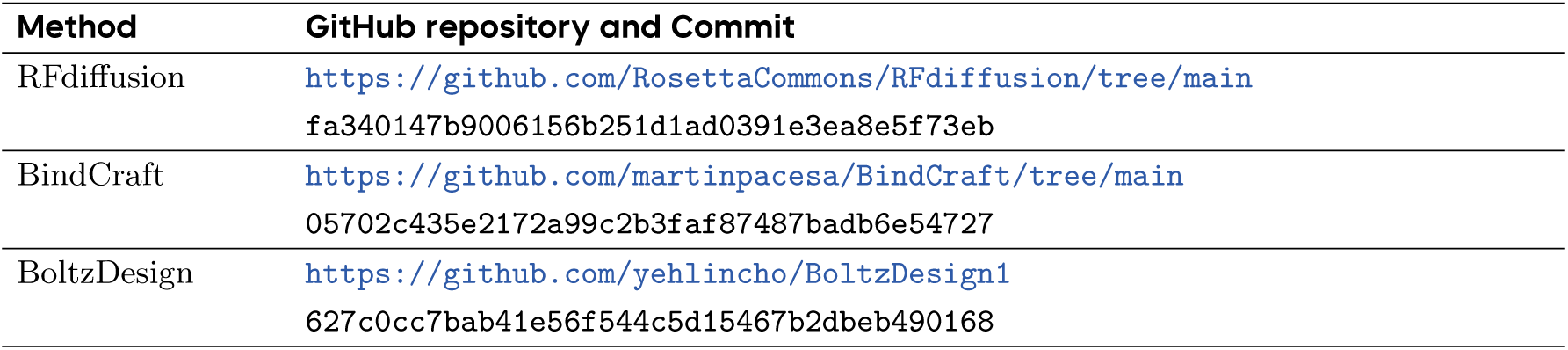
Details on running the compared methods.

**Table 5.**
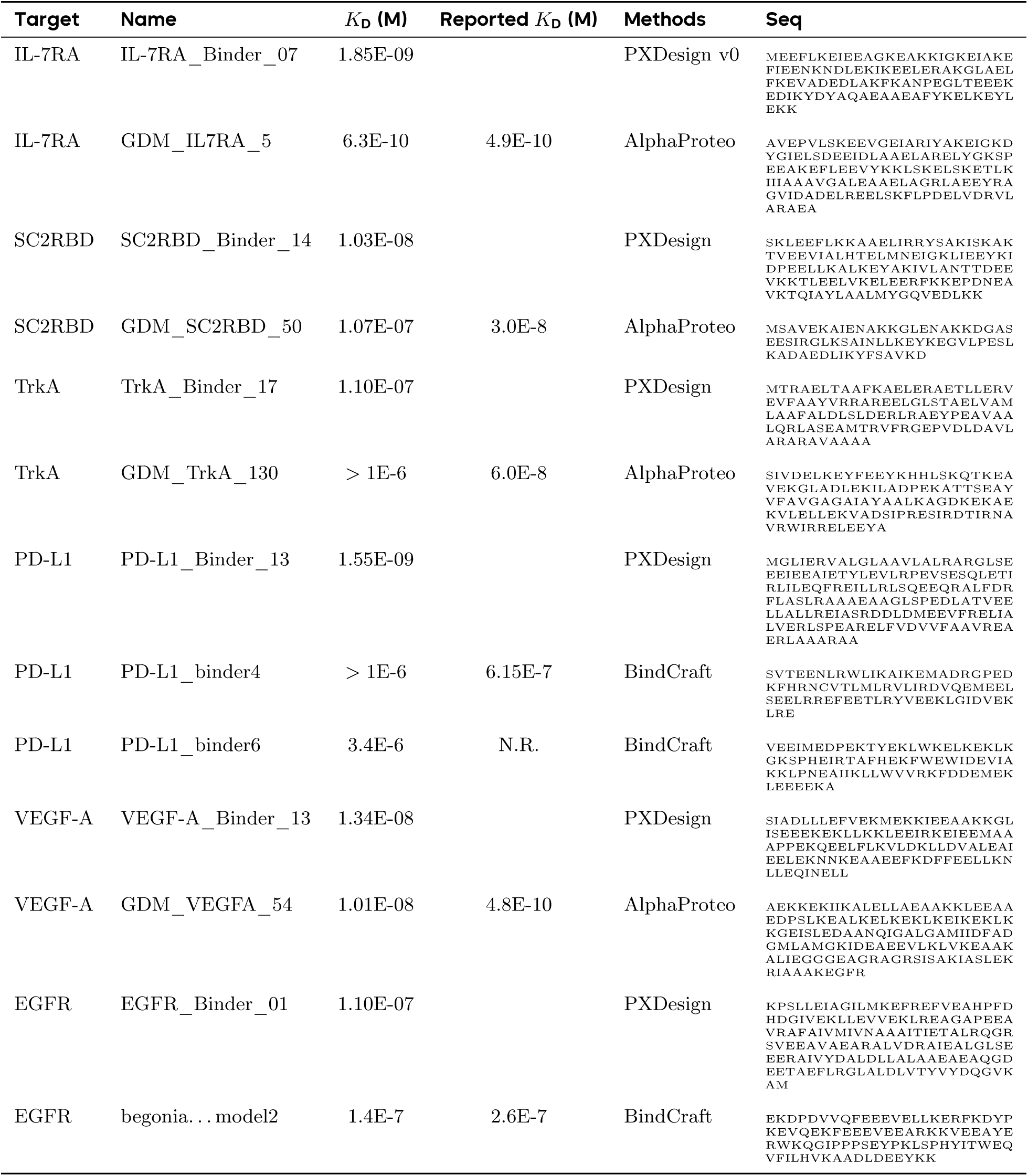
Sequences and binding affinities. *K*_D_ values are from our measurements, and “Reported” *K*_D_ values are taken from the original publications when available. Values prefixed by “*>*” indicate lower bounds; “N.R.” denotes values that were not reported.

### **E** Details of Wet-lab Results

1. **PXDesign**: We report the hit rate by classifying binders exhibiting a *K*_D_ lower than 1000 nM as hits. The hit rate is defined as the proportion of positive binders relative to the total number of designs sent for wet-lab characterization.
2. **AlphaProteo**: Table 1 of the main text reports success rates based on yeast display, which is known to have frequent false positives. We therefore corrected these rates using the true positive ratio from HTRF validation results (Table S3).
3. **Chai-1d / Chai-2**: Results were obtained by manual reading of Figure 2a and 2b.
4. **Latent-X**: For Latent-X, “detectable” corresponds to Tier 1 hits defined by an HT-BLI response *>* 0.03 RU, while “*K*_D_ measured” denotes designs with experimentally measured *K*_D_ *<* 1000 nM from Tier 2 assays.
5. **BindCraft**: For PD-L1, the experimental success rates were retrieved directly from Figure 1 of the BindCraft paper [38].
6. **Boltzgen**: We collected the experimental success rates for PD-L1 and IL-7RA from the preprint of Boltzgen [40].
7. **Proteinbase**: For EGFR, success rates were retrieved from ProteinBase (https://proteinbase.com) on Dec 8th 2025, filtering for mini-proteins designed using either RFdiffusion or BindCraft.
8. **RFdiffusion**: We manually extracted the success rates from Figure 6b of the RFdiffusion paper [3].

### **F** Cyclic Peptide Details

#### Benchmark Settings

**Target information.** We benchmarked the performance of PXDesign in cyclic peptide binder design against 12 diverse protein targets: 8 minibinder targets from AlphaProteo [59] (BHRF1, SC2RBD, PD-L1, TrkA, IL-7RA, IL17A, VEGF-A, TNF-*α*), 3 targets with experimentally determined structures from RFpeptides [42] (MCL1, MDM2, GABARAP), and 1 target from AfCycDesign [41] (Keap1). For the AlphaProteo and RFpeptides targets, we adopted the cropping schemes and hotspot selections described in the respective original studies. For Keap1, as AfCycDesign primarily employed hot-loop grafting rather than *de novo* design, we generated cyclic peptides using the structure with PDB ID 2FLU [33], applying a cropping range of A325–609 and hotspot residues A334, A380, A382, A415, A483, and A530, according to previous structural biology research [33].

**Method setup.** For PXDesign, we incorporated the Type 2 cyclic offset as described in AfCycDesign [41] into the positional encoding module on the cyclic chain. All other settings followed the minibinder design protocols detailed in Appendix B and Appendix C. For the baseline method, RFpeptides [42], we used the RFdiffusion release specified in Table 4. Each method was used to generate 128 peptide sequences of lengths 8–18, considering synthetic feasibility, developability, structural stability, and prior successful designs.

To assess real-world efficiency, we estimated the number of successful designs within 24 hours on an NVIDIA V100 GPU. The *in silico* success rate for cyclic peptide binders is determined using the AF2-IG-easy filtering criteria, encouraged by prior work leveraging AlphaFold2-based filters [41, 42]. As large-scale experimental binding data for cyclic peptide binders remain unavailable to our knowledge, we leave the validation and refinement of these *in silico* filters for future investigation.

### Additional Results

In addition to the overall *in silico* success rates for cyclic peptide binder design tasks shown in Figure 6a, we report detailed per-length success rates across different targets in Figure 12.

